## Supplemental Figures and Tables for "Assembly of a young vertebrate Y chromosome reveals convergent signatures of sex chromosome evolution"

**Figure S1**. Canu assembled contigs were aligned to the reference genome. Contigs that aligned to the autosomes form a clear unimodal distribution, whereas contigs aligned to the X chromosome do not. The increased number of contigs aligned to the X chromosome with lower sequence identity were separated as putative Y chromosome contigs.

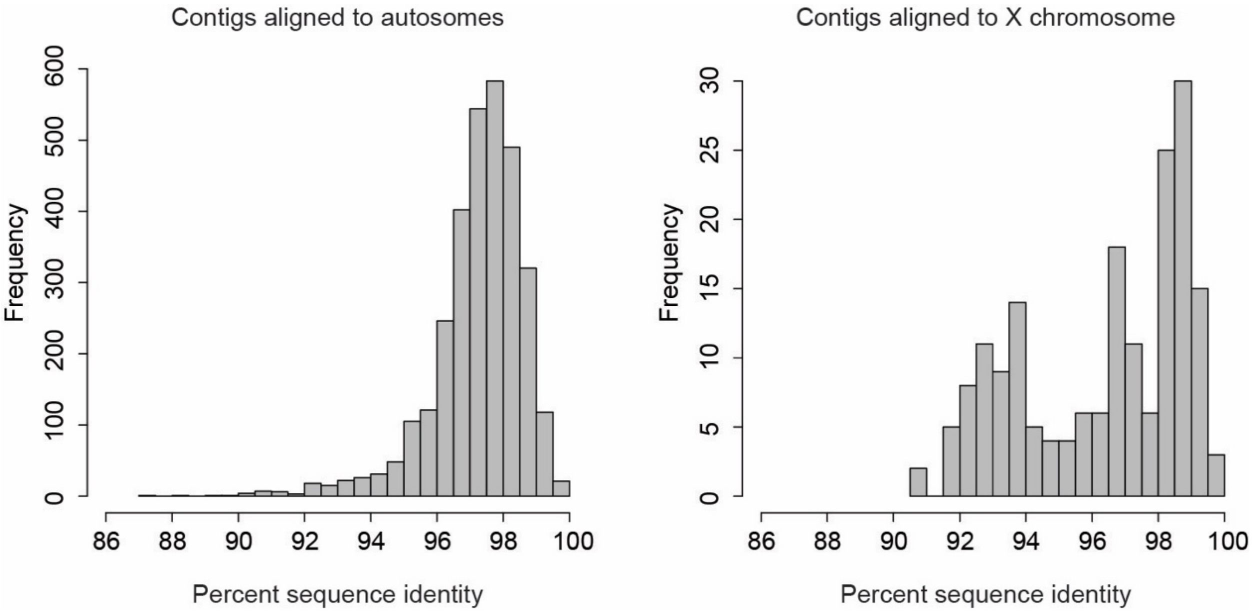

**Figure S2.** The PacBio assembly pipeline accurately reconstructed the X chromosome. The PacBio assembled X chromosome was split into three main scaffolds, with the two smallest scaffolds corresponding to the pseudoautosomal region and the third larger scaffold mostly aligning to the remainder of the reference X chromosome. Alignments between the PacBio assembled X chromosome and the reference X chromosome are largely colinear. Positions are shown in megabases.

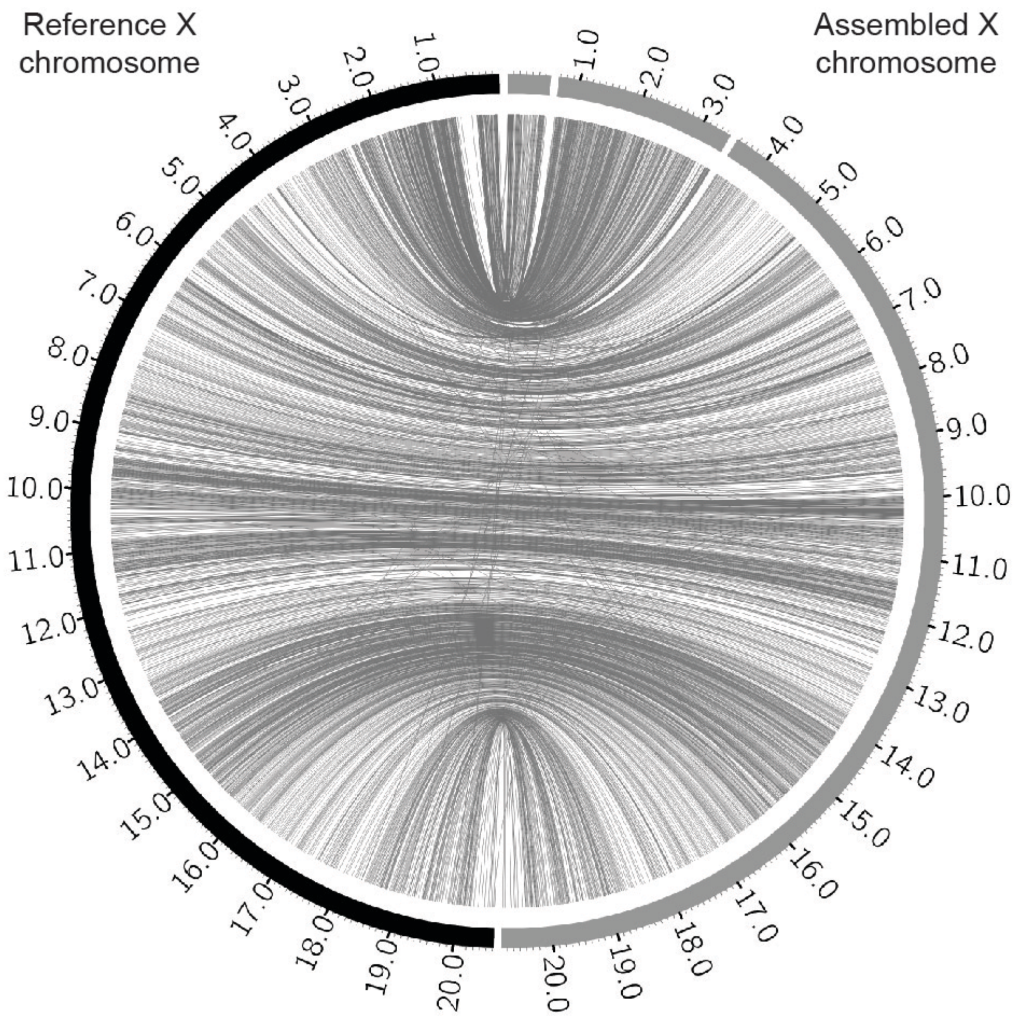

**Figure S3**. Hi-C chromosome conformation capture sequencing generated a single Y chromosome scaffold. The contact matrix shows a mis-joining of contigs at one end of the scaffold that includes fewer short-range interactions at the diagonal and an absence of long-range interactions elsewhere in the chromosome off the diagonal (upper left of diagonal). These contigs were removed for further analysis. Contig boundaries in the assembly are denoted by the black triangles along the diagonal.

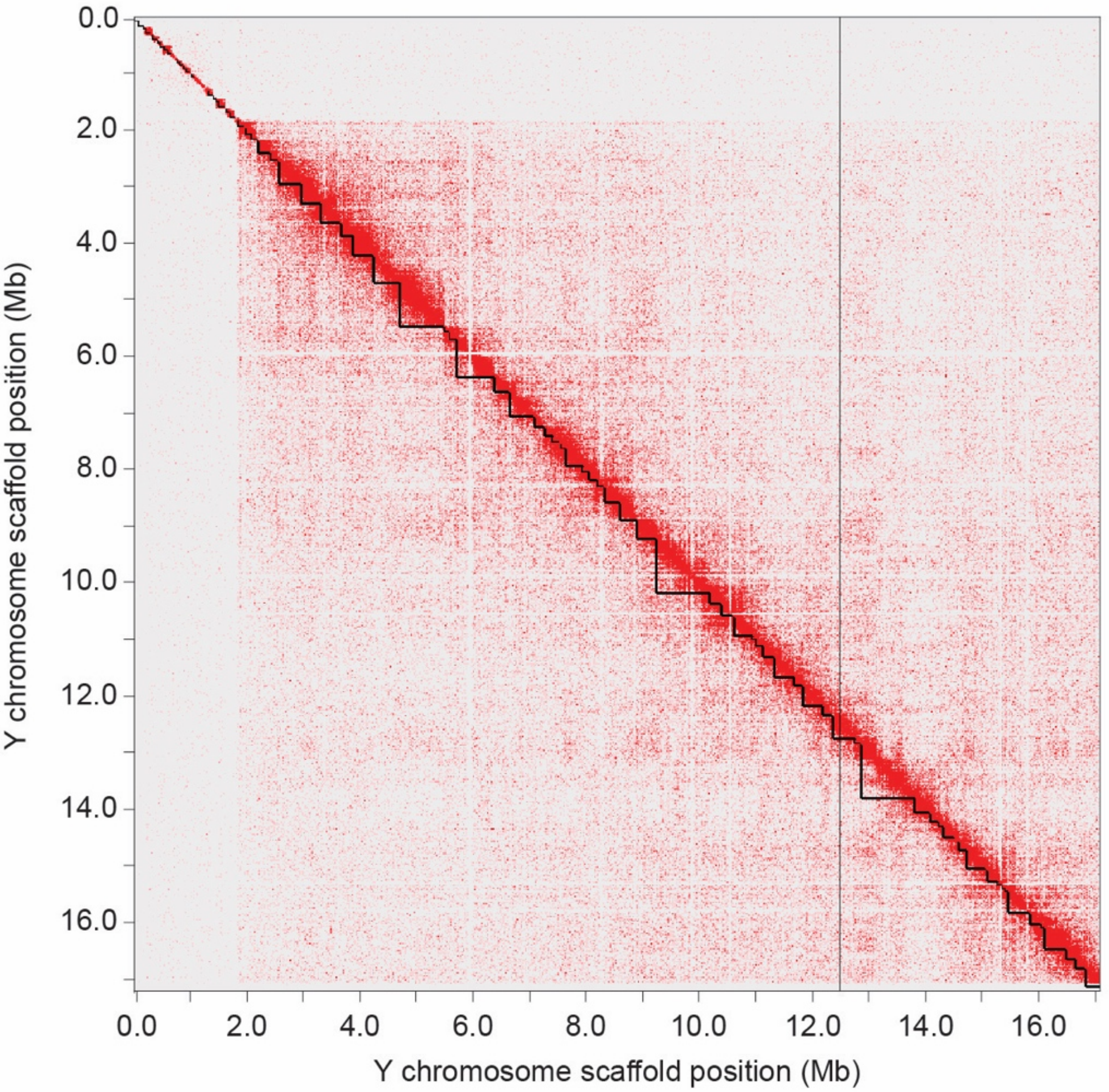

**Figure S4**. Stratum one is located on the end of the Y chromosome, opposite of *Idh*. Fluorescent *in situ* hybridization probes were generated from a stratum one BAC (CHORI 215-013C20). The BAC (pink) is located at the end of the Y chromosome (arrow), opposite from *Idh* (green) on mitotic metaphase spreads. *Idh* is located centrally on the X chromosome (arrowhead).

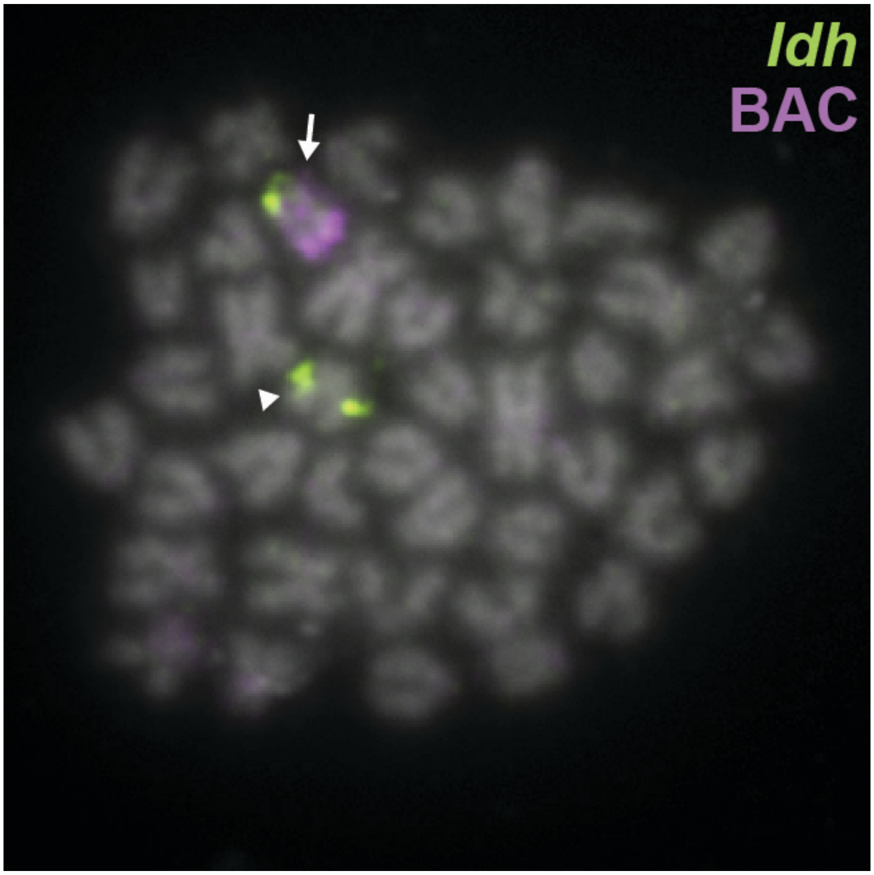

**Figure S5.** Short-read sequences from a chromatin-immunoprecipitation (ChIP-seq) with CENP-A from a second male fish were aligned to the reference Y chromosome assembly. There is a prominent peak between markers STN187 and WT1A where the centromere is located in cytogenetic maps.

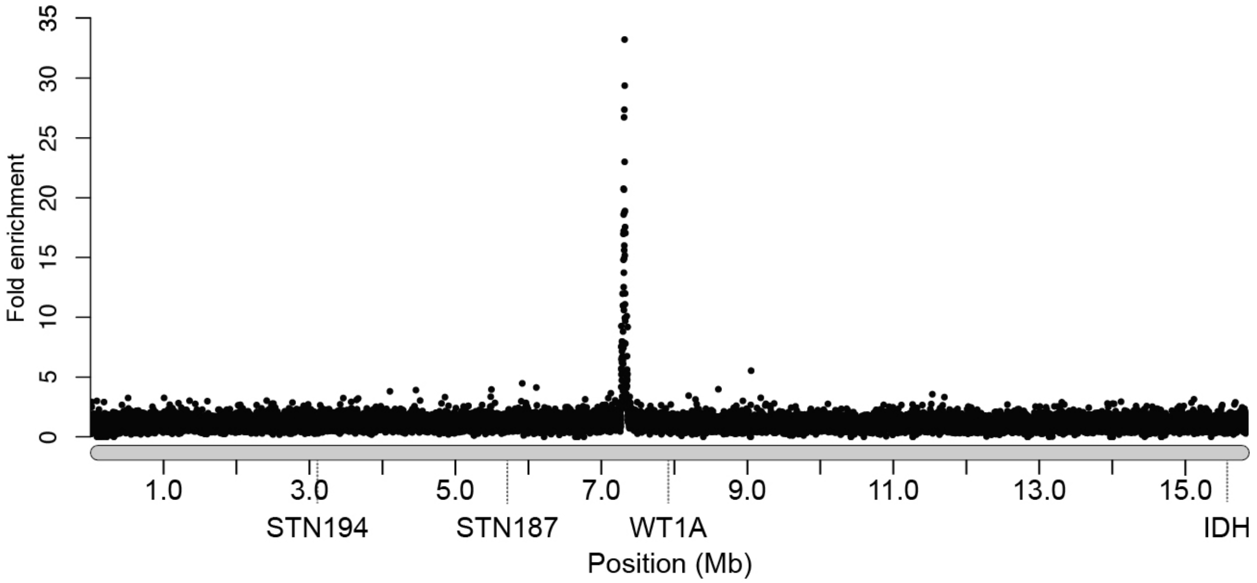

**Figure S6.** Alpha satellite monomeric repeats found on the Y chromosome show conservation with the core centromeric repeat found on the autosomes and the X chromosome. The putative CENP-B Box is shown in red.

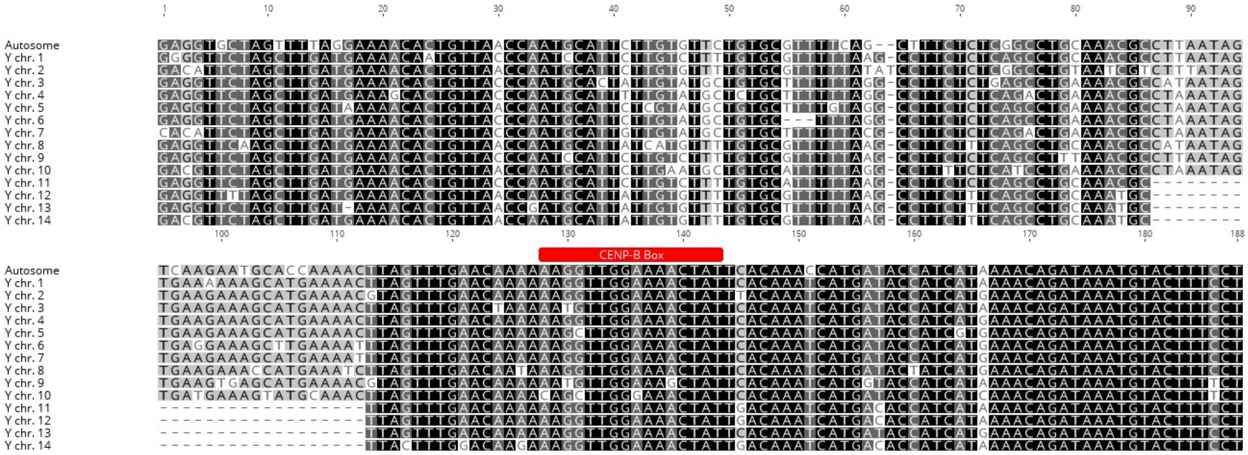

**Figure S7.** Transposable elements are at a higher density across the Y chromosome relative to the X chromosome. Repeat families were identified using a combination of RepeatModeler and RepeatMasker. The total number of repeats across 250 kb bins are shown.

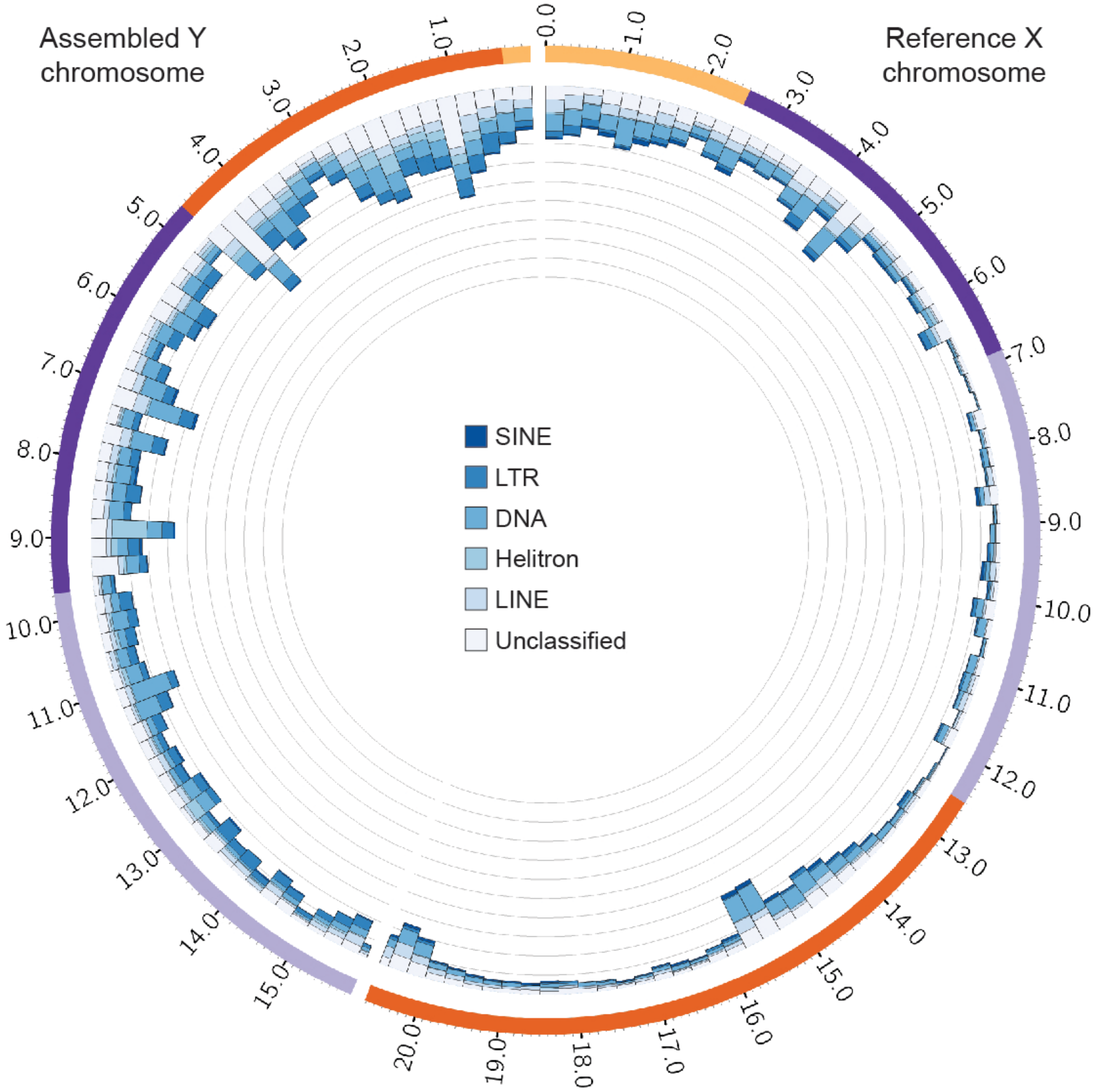

**Figure S8.** Alignment of protein coding sequences of vertebrate *Amh* genes. Conserved residues across taxa are highlighted in black. The functional protein Amh and TGF-β are shown in red.

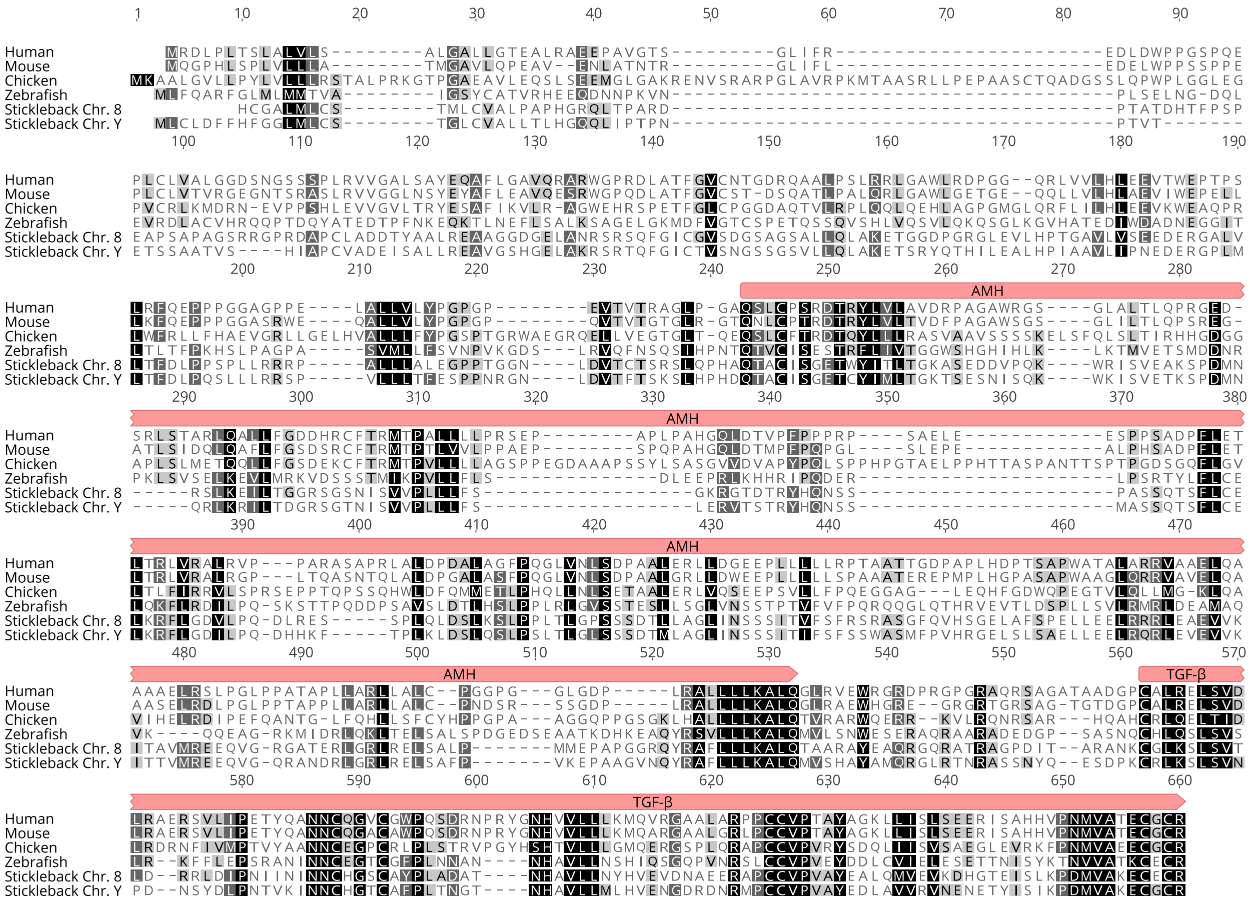

**Figure S9.** A putative model for the evolution of the Y chromosome indicating there were four separate inversions. Inversions are shown by dotted lines. Cytogenetic markers that could not be detected on the Y chromosome are shown degraded by lighter colors. The centromere position is indicated between markers WT1A and STN187.

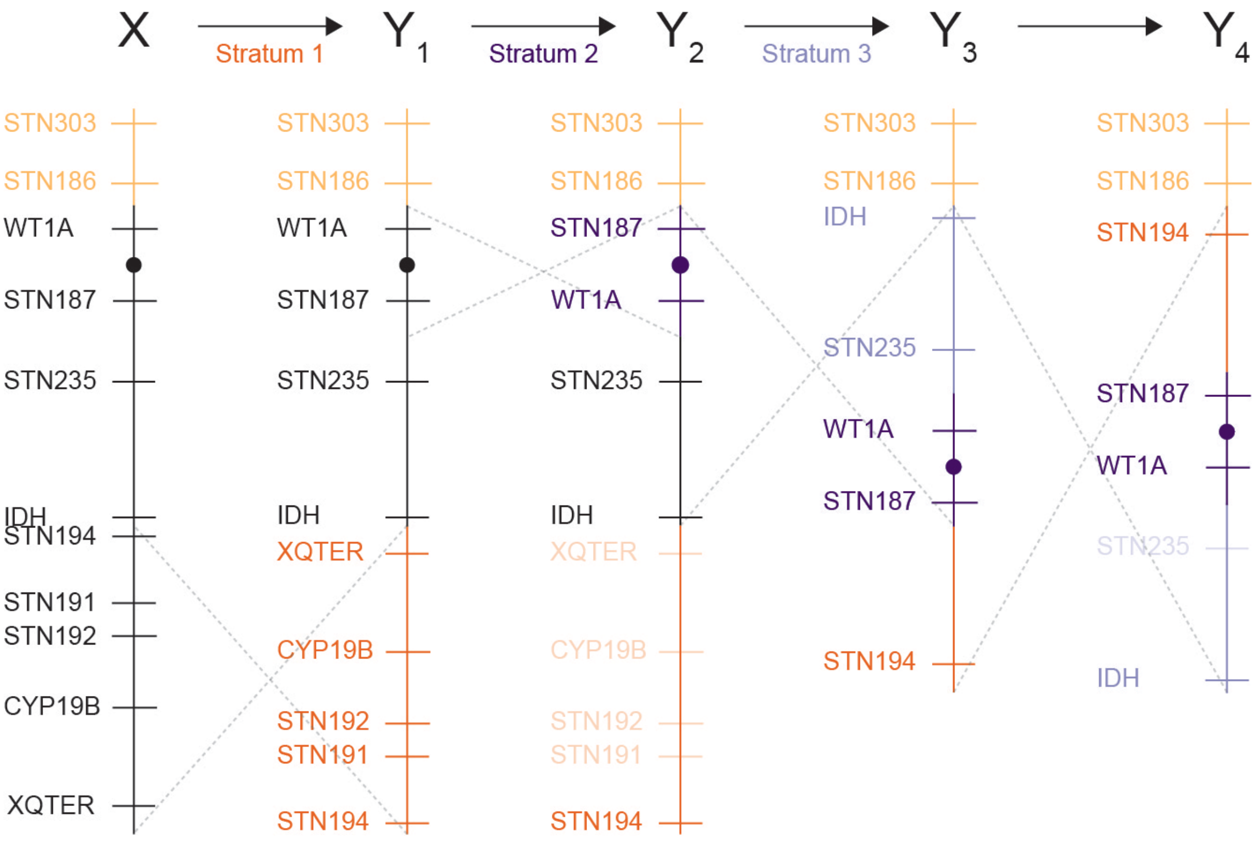

**Supplemental table 1**. Sequencing statistics before and after assembly with Canu.

| | PacBio preassembled reads | Canu assembly |
|---|---|---|
| **Total sequence length** | 34.84 Gb | 622.30 Mb |
| **Total contigs** | 3,255,924 | 3593 |
| **Contig N50 length** | 21,068 bp | 541,786 bp |
| **Max. contig length** | 87,617 bp | 12,930,346 bp |
| **Min. contig length** | 1000 bp | 1028 bp |
| **Mean contig lenth** | 10,701 bp | 173,198 bp |
| **Median contig length** | 5969 bp | 56,654 bp |

**Supplemental Table 2.** $Log_2$ fold change between testis tissue and three other tissues for genes that are duplicated on the Y chromosome and have an X-linked homolog.

| Comparison | Y duplicated | X homolog | P value |
|---|---|---|---|
| **Testis versus Liver** | -1.300 | -0.186 | 0.025 |
| **Testis versus Brain** | -3.379 | -0.114 | 0.006 |
| **Testis versus Larvae** | -5.395 | -4.872 | 0.998 |

**Supplemental Table 3**. The effect of sequence identity threshold on the total number of X chromosome contigs identified.

| Sequence identity threshold | Number of X chr. contigs | Total contig length | Difference from reference X chr. |
|---:|---:|---:|---:|
| 94% | 133 | 23,137,131 bp | 2,518,665 bp |
| 95% | 124 | 21,909,126 bp | 1,290,660 bp |
| 96% | 114 | 21,255,474 bp | 637,008 bp |
| 97% | 90 | 18,109,484 bp | -2,508,982 bp |
| 98% | 73 | 15,892,396 bp | -4,726,070 bp |

**Supplemental Table 4.** The effect of the 3D-DNA parameter --editor-repeat-coverage on the number of concordant BACs that align to the assembly.

| Editor repeat coverage | Number of concordant BACs |
|---|---|
| 8 | 62 |
| 9 | 48 |
| 10 | 89 |
| 11 | 92 |
| 12 | 92 |
| 13 | 92 |
| 14 | 89 |
| 15 | 89 |
| 16 | 89 |
| 17 | 88 |
| 18 | 88 |
